# Degradation of fish food webs in the anthropocene

**DOI:** 10.1101/2024.10.22.619402

**Authors:** Juan D. Carvajal-Quintero, Maria Dornelas, Lise Comte, Juliana Herrrera-Pérez, Pablo A. Tedesco, Xingli Giam, Ulrich Brose, Jonathan M. Chase

**Author notes:** Corresponding author: Juan D. Carvajal-Quintero (;). Co-senior authors (Ulrich Brose, Jonathan M. Chase).

## Abstract

The Anthropocene is marked by profound changes in biodiversity and the ecosystems in which species live ^1–4^. A primary signature of this change is the often rapid change in species composition through time (i.e., species turnover) rather than changes in the numbers of species per se ^5–7^. Less well known, however, is which types of species are ‘winning’ and which are ‘losing’ as ecosystems change through time ^8,9^, as well as whether and how these changes influence higher-level processes in the food webs in which species are embedded ^10,11^. Here, we combine a compilation of long-term observations of ∼15,000 freshwater and marine fish communities surveyed for 1949-2019 years, together with information about their diets and trophic status in order to evaluate how the food webs in which these fish communities are embedded are changing through time. We found widespread alteration to fish food web topology and functioning. This includes an increase in connectance and generalism in food webs, which has led to greater predation pressure, as indicated by higher diet overlap and increased prey vulnerability. We also identified a decrease in modularity, which has reduced the compartmentalization within local networks. These changes extend across the trophic structure of food webs, causing a cascading shift in the proportion of species across trophic levels. Our study highlights the complex responses of biodiversity change of fish food webs in the Anthropocene, which can ultimately influence the functions of these ecosystems and human well-being ^12,13^.

## Main

Anthropogenic activities have extensively impacted nature and the biodiversity within, leading to rapid changes in the numbers and types of species in local assemblages ^3,5,6,14–16^. These changes often favor species with certain traits and disfavor others ^17–21^, which can in turn alter patterns of ecosystem functioning and related ecological processes ^22–24^. For example, anthropogenic activities often disproportionately alter the body size of species within populations and assemblages ^18,25–27^, and these changes can transform the structure and function of food webs ^28–32^. However, because trends in body size vary substantially among communities ^18^, it is not yet clear what, if any, are the consequences of these changes to the structure and functioning of food webs.

Conceptually, cumulative shifts in species composition turnover and body size changes can alter the structure and function of food webs via several alternative pathways (Fig. 1). First, there can be no change in local species richness from the community at time 0 (Fig. 1 T0), as is often observed ^5–7^. If there is no change in species richness, and also no change in important traits like body size (Fig. 1 scenario T1A), we would expect no change in food web structure.

**Fig. 1.**
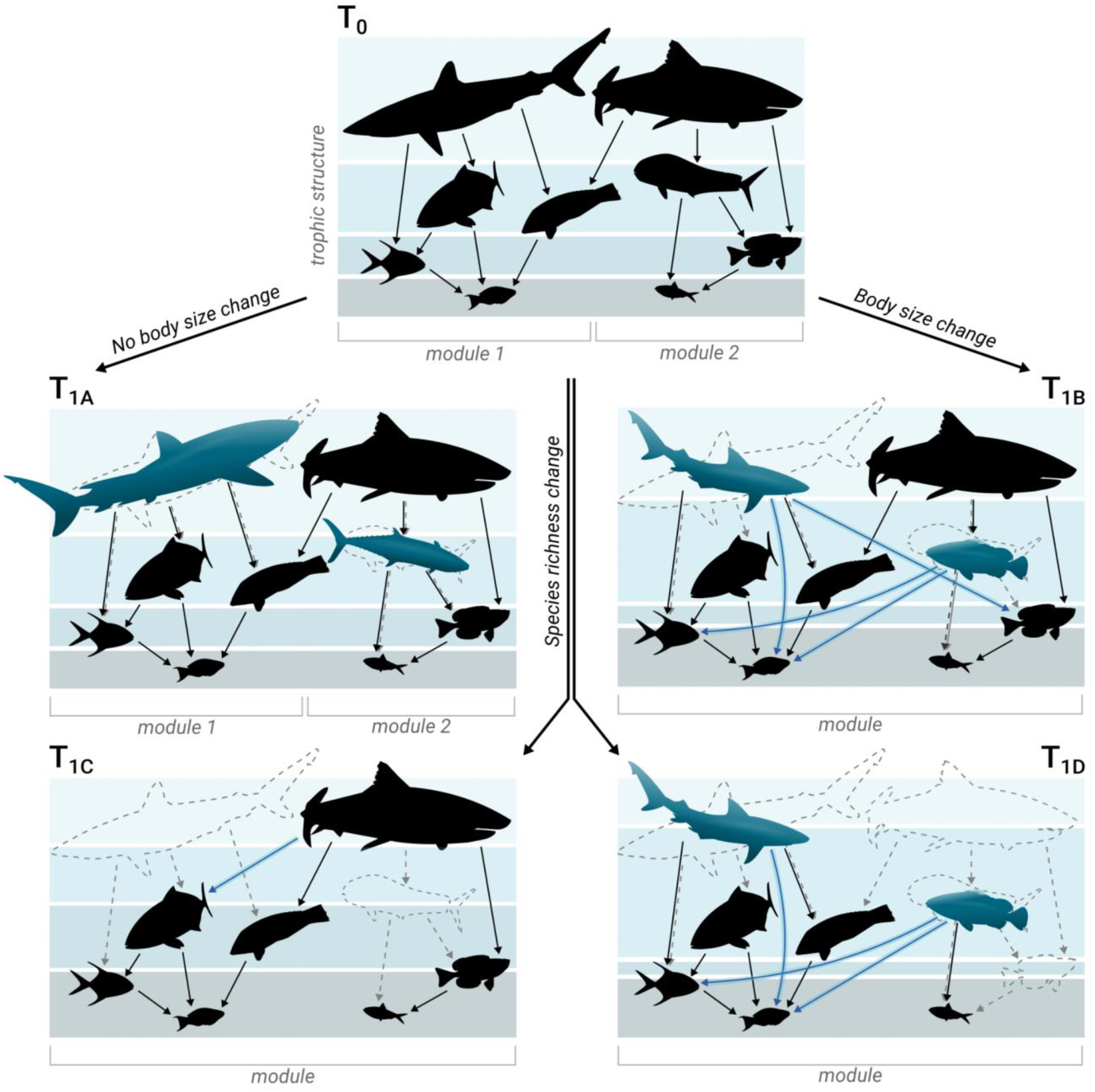
Pathways of temporal changes in taxonomic diversity and body size resulting in different potential scenarios of food web changes. Species turnover (represented by species dashed and blue silhouettes) unfolds from T_0_ (top) to T_1_ (below), reshaping species composition and potentially impacting species richness and body size. Changes in species composition not affecting body size are depicted on the left (T_1A_ and T_1C_), while those altering body size on the right (T_1B_, T_1D_). When turnover does not affect species richness nor body size, no structural changes are expected in the food webs (T1A). However, alterations in body size arrangement can change the prey selectivity of the species (blue lines) inducing food web shifts even in the absence of changes in species richness (T1B). When species turnover alters species richness, it directly impacts food web topology by changing the number of interacting species and links (T1C). Moreover, if turnover reduces both species richness and body size, cumulative effects on food web structure and functioning are anticipated (T1D).

However, even with no change in species richness, the average body size of species in the community can decrease because of changes in community composition, as sometimes observed (Fig. 1 scenario T1B). Here, we would expect a shift in the structure and function of the food web. This is because the trophic niche of a species is often determined by its body size.

Therefore, the selective filtering of body sizes can introduce novel trophic interactions, rewiring the food webs by reshaping the density and distribution of links, and consequently altering their structure and functioning ^28,30,32,33^. If species richness changes through time, another set of possibilities can emerge. For brevity, we consider scenarios where species richness declines without (Fig. 1 scenario T1C) or with (Fig. 1 scenario T1D) concomitant changes in body size, but species richness increases would simply be the inverse. If species richness declines, but body size does not change (Fig. 1 scenario T1C), we expect a change in the structure of the food web via changes in the number of nodes in the food web, and consequently a less connected network (lower connectance) due to the decrease in the number of interactions ^34,35^. Alternatively, if both species richness (number of nodes) and body size change (Fig. 1 scenario T1D), we would expect a simultaneous reduction in the network size and connectance along with the food web rewiring caused by the changes of predator-prey interactions resulting from alterations in species’ body sizes ^28,30,32,33^.

Here, we present a comprehensive assessment for how the alteration of species composition and body size have reshaped fish food webs in the recent past. To do so, we compiled an extensive database of fish assemblage time series from freshwater and marine ecosystems (RivFishTIME and BioTIME ^36,37^). These data encompass a diverse array of 15,029 fish assemblages containing 2,844 fish species, and were sourced from assemblages surveyed in 103 studies from across the world (Extended Data Fig. S1). Time series ranged from 2 to more than 90 years. After standardizing sampling effort across time series ^5,38^, we assessed temporal trends in species richness, composition dissimilarity, and body size. We then linked these shifts in the taxonomic and functional layers to changes in reconstructed food web metrics (e.g., network connectance and modularity) that are associated with food web stability and resilience ^39–41^ using a predator-prey model calibrated with a comprehensive database of over 23,000 trophic interactions and body size records for co-occurring fish species pairs (see Methods in Extended Data). To explore temporal trends in these metrics, we employed hierarchical generalized linear models, nesting spatial-unit time series within their original studies to address potential biases arising from the non-independence of spatial-unit time series within a given study (further details are provided in the Methods Extended Data). We fit each biodiversity metric (i.e., taxonomic diversity, body size and food web metrics) separately, and reported sensitivity of the rate of changes (model slope) to the replicate variability. We considered any 95% confidence interval (CI) not overlapping zero as compelling evidence for a directional trend (see details in Methods).

We found no overall trend in species richness through time (mean: 0.0001; 95% CI: −0.0003, 0.0006; Fig. 2) but a strong directional change in species composition (mean: 0.0047; 95% CI: 0.0040, 0.0055; Fig. 2), an increasingly well-known phenomenon across taxa ^5–7,42^. More importantly, we found that these compositional changes were nonrandom, with selective filtering towards smaller species reflecting an overall downsizing of body size in species assemblages over time (mean: −0.0009; 95% CI: −0.0012, −0.0006; Fig. 2). This downsizing of fish communities has been observed elsewhere ^18^ and is frequently associated with human-driven overexploitation, warming, and reduced resource availability ^25,43,44^. Perhaps more importantly, however, owing to the pervasive importance of body size ^32,45^, changes in assemblage-level body sizes are likely to alter the structure of food webs and the ecosystems in which these changes take place.

**Fig. 2.**
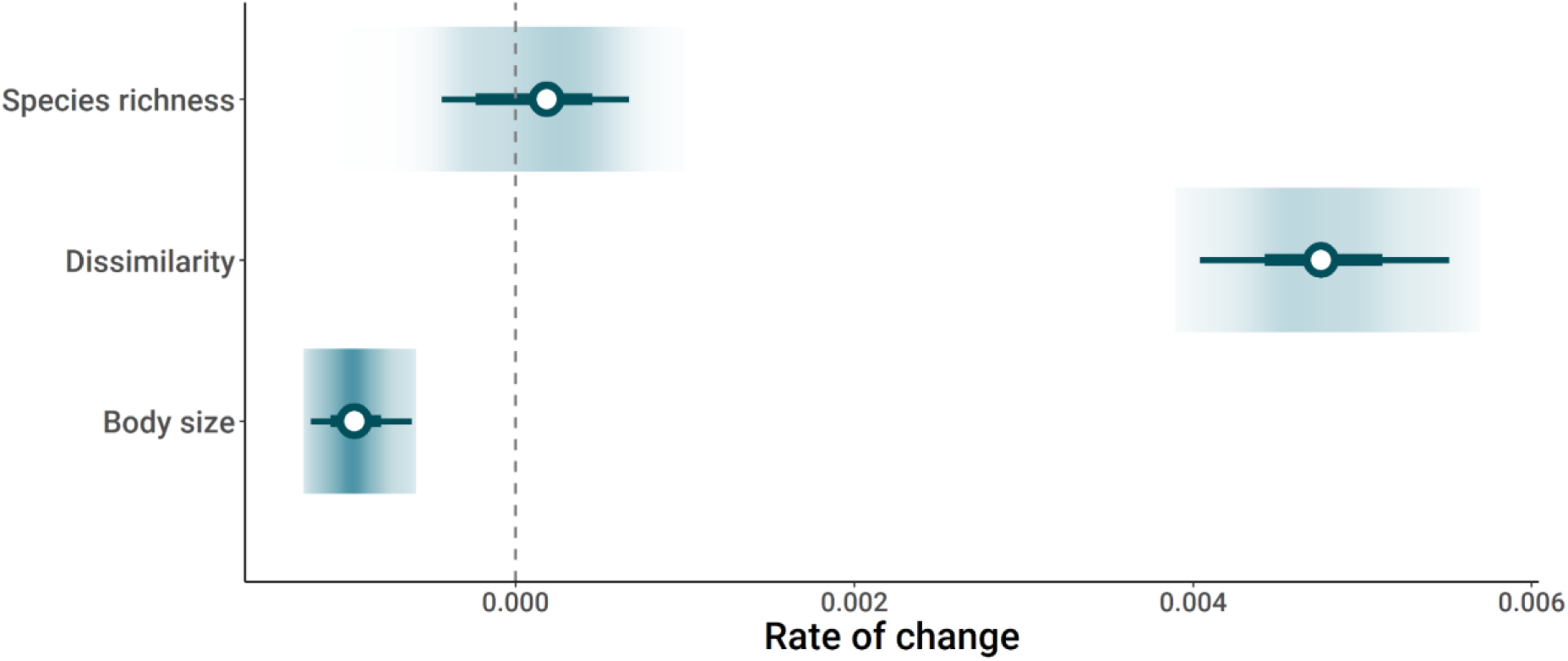
Changes in assemblage metrics. The gradient plots display the distribution of slopes for changes in species richness, dissimilarity, and body size. Darker colors correspond to higher densities. The horizontal bars with error bars denote the mean and the 50% and 95% confidence intervals (CIs) of the mean estimates (depicted by white circles).

### Topological and functional changes in food webs

Over time, we found that the connectance of fish food webs increased (mean: 0.0007; 95% CI: 0.0004, 0.0010; Fig. 3). This was because there was an increase in mean network generality (mean: 0.0199; 95% CI: 0.0083, 0.0300; Fig. 3), indicating that temporal turnover favors more generalist species that feed on a broader range of prey. Such replacement of specialists by generalists is a frequently observed signature of perturbed environments ^17^, and thus we expect increases in network connectance to be a general phenomenon. We also observed an overall decrease in food web modularity through time (mean: −0.0010; 95% CI: −0.0014, −0.0007; Fig. 3), which describes the degree of species frequently interacting in clusters. This decline in modularity can be attributed to two processes. First, modularity can decline through time because of increases of generalist species, which disrupt the typical block-like structure of ecological networks by interacting broadly with a diverse prey array ^10,46^. Second, modularity can decline because of a reduction in the proportion of top predators (mean: −0.1414; 95% CI: −0.1505, −0.1316; Fig. 4) that anchor modules in ecological networks by interacting with various species at lower trophic levels, forming distinct subgroups of interactions ^12,32,47,48^. These topological changes may have important implications for the stability and the maintenance of biodiversity, since species interaction networks largely influence the response of community and ecosystems to environmental change ^2,49^. For example, the increased connectance and reduced modularity can destabilize ecosystems by synchronizing the responses of food webs to perturbations ^10^.

**Fig. 3.**
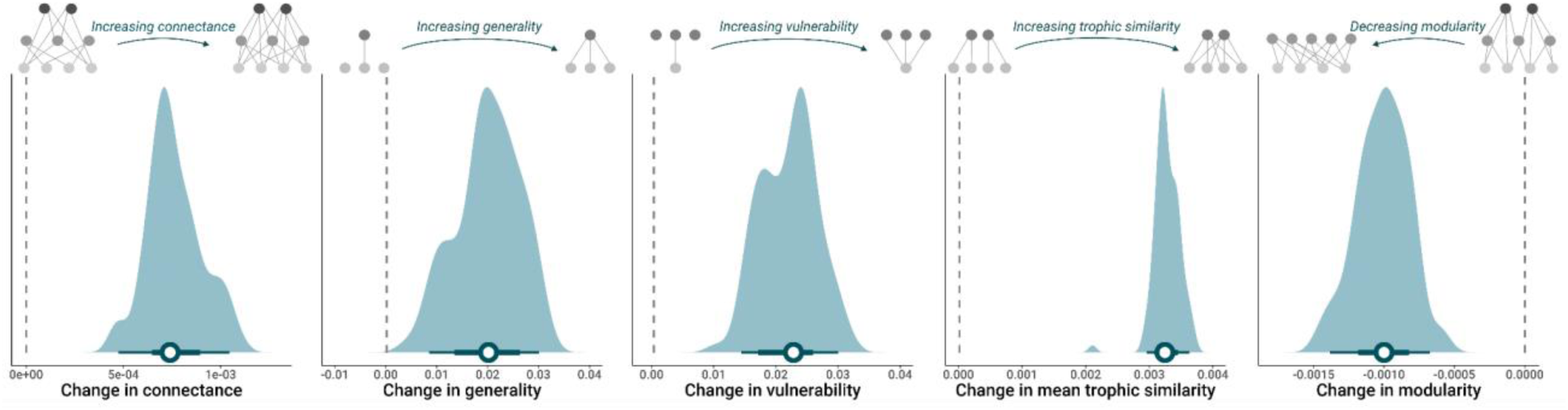
Changes in food web topology metrics and trophic similarity. The density plots display the distribution of slopes for changes in connectance, generality, vulnerability (predation pressure), trophic similarity, and modularity. The horizontal bars with error bars denote the mean and the 50% and 95% confidence intervals (CIs) of the mean estimates (depicted by white circles). The top insets illustrate the changes in the number and distribution of links that each trend represents, with the arrow representing the direction of the change from T0 (aligned with the dotted line representing no change) to T1.

**Fig. 4.**
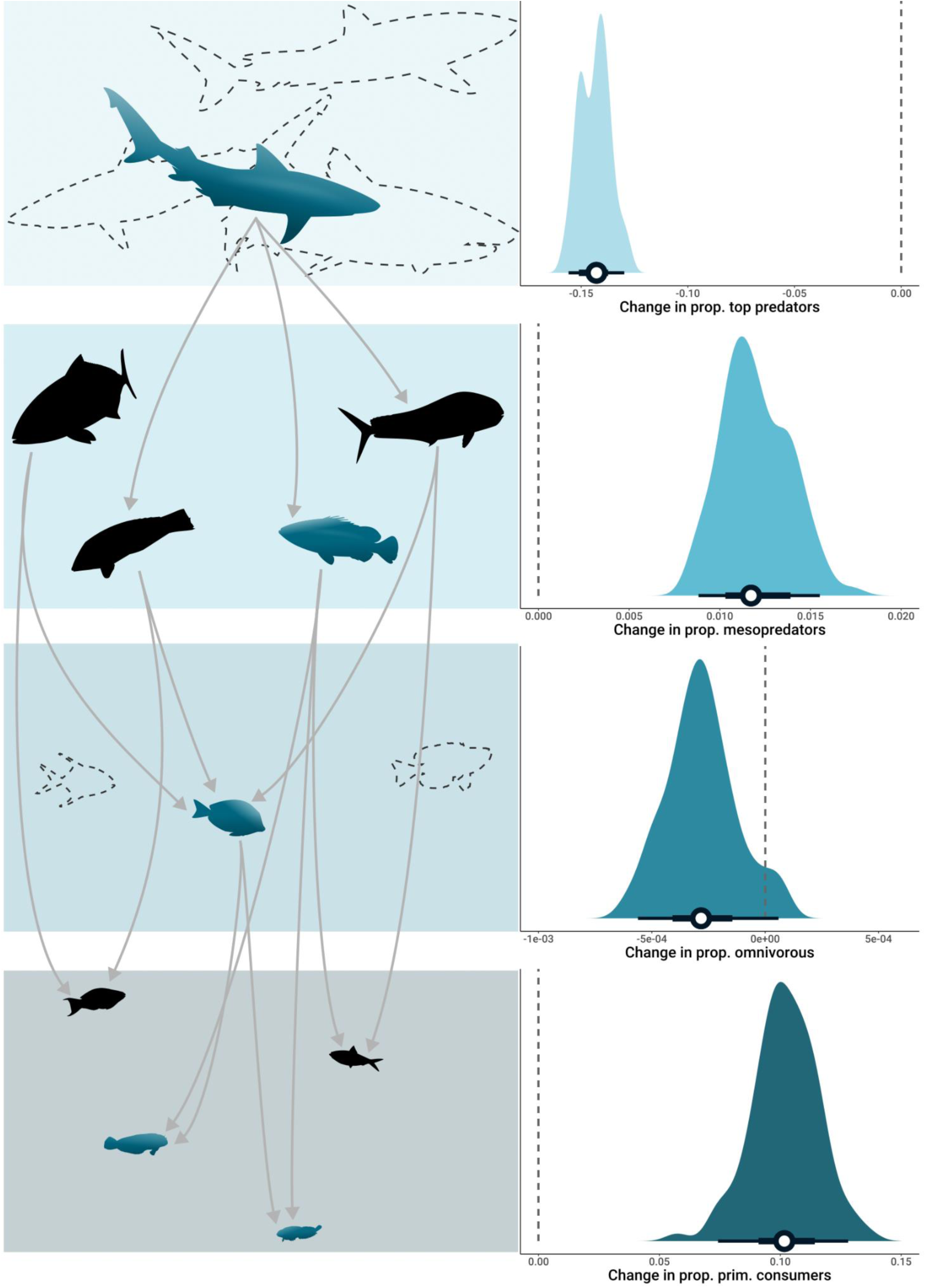
Changes in proportion of trophic groups. The density plots (right) display the distribution of slopes for changes in proportion within the assemblages of top predators, mesopredators, omnivorous, and primary consumers. The horizontal bars with error bars denote the mean and the 50% and 95% confidence intervals (CIs) of the mean estimates (depicted by white circles). On the left are illustrated schematic changes in the proportion of species across trophic groups to help with the interpretation. Blue silhouettes represent new species, dashed silhouettes indicate species lost, and black silhouettes denote species present across the entire food web time series.

Reductions in the degree of modularity can also compromise the stability of food webs because it fosters frequent interactions within clusters that previously limit the propagation of species-specific impacts across the food web ^39,40,50,51^.

At the functional level, we found increases in mean network generality (mean: 0.0199; 95% CI: 0.0083, 0.0300; Fig. 3), trophic similarity (mean: 0.0032; 95% CI: 0.0029, 0.0035; Fig. 3), and prey vulnerability (mean: 0.0217; 95% CI: 0.0137, 0.0295; Fig. 3). These indicate that temporal turnover favors generalist species —those with wider trophic habits— thereby increasing diet overlap and predation pressure, as prey are targeted by more predators. Indeed, generalist predators tend to be more adaptable to changing environmental conditions, enabling them to colonize new assemblages ^10,17,52^, which prompts local extinctions of specialists, making food webs more susceptible to environmental disturbances ^10,31^. These processes can ultimately lead to the overexploitation and reduction of prey species, triggering cascades of extinctions ^53^.

Across all time series, we found cascading shifts among adjacent groups in the trophic hierarchy. Specifically, the proportion of species of top predators and omnivores within the food web declined, while mesopredators and primary consumers increased through time (Fig. 4). This pattern of opposite trends among adjacent trophic groups emphasizes the interconnected nature of the trophic hierarchy in food webs and the potential repercussions of altering one level on the entire food web. While trophic structures have tended to be stable for hundreds to many thousands of years amidst substantial shifts in species composition ^54,55^, our observation of widespread transformation in the trophic structure of aquatic food webs underscores the severity of Anthropocene impacts on biodiversity.

We found no systematic differences in our results when we repeated our analysis with time series spanning different time periods (Extended Data Fig. S3 and Fig. 4), suggesting these findings are robust to time series length. In addition, the trends we observed in food web structural changes were similar across both freshwater and marine environments (Extended Data Fig. S5 and Fig. 6), indicating that the degradation of food webs is widespread across ecosystems. Nevertheless, we acknowledge the limitations stemming from the geographical biases of the data used in our study. For example, like most such syntheses, our data are notably biased towards Europe, North America, and Australasia (Extended Data Fig. S1). The challenge of accurately quantifying biodiversity trends is heightened by the scarcity of long-term biodiversity data in many parts of the world, particularly in tropical regions (e.g. Achieng *et al.* 2023). Thus, our estimated mean trends are not suitable for geographical extrapolation, but rather can only serve as a representation of the most current state of knowledge derived from existing open data ^57,58^.

In all, we show that selective turnover of species via reductions in body size is degrading the topology and trophic structure of fish food webs in the Anthropocene. Notably, our findings emphasize that, despite the absence of a general trend in species richness change over time ^5–7,42^, a noticeable transformation in the functioning of biodiversity within food webs is occurring through the filtering of new trait arrangements. Food webs are becoming structurally less complex due to the loss of modularity, while becoming more generalized with increases in connectance, diet breadth and overlap. While we can only speculate on the causes of these fish food web changes across many different systems experiencing an array of different human threats, we suspect they are tied to the historical and ongoing degradation of fish fauna. Factors such as widespread and targeted exploitation of marine and freshwater biota, habitat destruction, and climate shifts have resulted in regional and local extinctions and population declines of some species, particularly larger predatory fish ^59,60^. Fish are crucial for supporting aquatic biodiversity, economies, livelihoods, nutritional security, and cultural practices on our planet ^61,62^, and as such, understanding the magnitude and timing of food web changes resulting from strong human-induced disruption is essential for their effective conservation and sustainable management ^41,63,64^. Our results highlight the importance of monitoring species interaction networks to better understand the functional effects of biodiversity change in the Anthropocene.

## Methods

Our analysis integrates ecological assemblage, trait data (body size), and documented trophic interactions to reconstruct time series of fish food webs, and elucidate the topological and functional changes occurring within them. We drew upon two existing sources of published ecological assemblages, extracting time series information for both freshwater and marine environments. Species’ body sizes and trophic interactions were sourced from a combination of published literature and open-access databases. Below, we outline the standardization process and the calculations employed for reconstructing the time series of food webs. Additionally, we present the statistical analyses detailing the trends observed in fish food webs during the Anthropocene era.

To ensure consistency in the merging of the different datasets and databases, we harmonized taxonomy across the databases following the methodology outlined by Grenié ^65^, utilizing FishBase ^66^ as a taxonomic reference to rectify synonyms and correct misspellings for fish using the ‘rfishbase’ package ^67^.

### Data collection and selection

#### Assemblage time-series data

We obtained ecological assemblage data from the BioTIME ^37^ and RivFishTIME ^36^ databases, which are the most extensive global, open-access repositories of assemblage composition time series. BioTIME encompasses studies on various taxonomic groups, whereas RivFishTIME specifically focuses on freshwater fish. We gathered the studies on both freshwater and marine fish from BioTIME and integrated them with those in RivFishTIME. Each study encompasses distinct samples surveyed consistently over time using a standardized methodology, in which the number of years sampled can change.

Due to variations in spatial extent across studies, we adopted established methods to standardize the spatial extent among studies, by pooling those studies that had large extents and multiple sampling locations into spatial units ^38,68^. For marine environments, we employed a global grid comprised of 96 km² hexagonal cells as spatial unit (utilizing dggridR ^69^), whereas for freshwater environments, we utilized a global layer of subbasins spanning on average 99 km² (based on HydroBASIN level 12, ^70^). Studies that were contained within a single spatial unit (i.e., grid cell or subbasin) remained unpartitioned. Subsequently, each sample received a unique combination of study ID and spatial unit based on its latitude and longitude. This assignment generated a distinctive identifier for each assemblage time series within spatial units, ensuring the preservation of both study and sample integrity.

To minimize the impact of unobserved species on our biodiversity change estimates, we computed the abundance-based coverage ^71^ for each (annual) sample within each spatial-unit-level time series. We excluded all samples with coverage less than 0.85 based on the ratio of detected to expected species richness ^38,68^. Furthermore, we retained only those time series that contained records of abundance.

Finally, we implemented a sample-based rarefaction to standardize the number of samples across years within each time series ^38,68,72^. In this process we prioritized surveys conducted in the same climatic season, distinguishing between the warm season (from April to September) and the cold season (from October to March) in the Northern Hemisphere, and vice versa for the Southern Hemisphere. The rarefaction was repeated 100 times for each time series, and the outcomes of each iteration were preserved to reconstruct the food webs.

#### Body size data

We retrieved body size trait data, specifically the maximum reported body length, from both FishBase, and published literature sources (see Supplementary Data).

#### Species diets

To compile species diet data from scientific literature, we gathered information from various sources. We collected diet data from FishBase ^66^ and the Global Biotic Interaction database (GloBI). Additionally, we curated an independent dataset of fish diet records by consolidating information from journal articles and natural history handbooks (see Supplementary Data). For journal articles, we conducted Google Scholar queries using the scientific names of each fish species to identify relevant studies and acquired records not included in FishBase and GloBI. In each diet record, we documented consumer-resource interactions at the species level for the consumer and, when possible, at the highest taxonomic level for the resource. In cases where taxonomic information was unavailable for the resources, we retained the common names (refer to Supplementary Data for details). The diet information was subsequently used to calculate the trophic levels of the species and to calibrate the niche model used to reconstruct the food webs (see below).

#### Trophic groups

Species were grouped into distinct trophic groups based on their trophic levels (TL) using predefined thresholds from FishBase ^66^. The trophic groups include top predators (TL > 4), mesopredators (TL = 2.8 – 4), omnivores (TL = 2.2 – 2.79), and primary consumers (TL = 2 – 2.19). The calculation of TL values was performed using the ‘dietr’ package ^74^. This methodology calculates fractional trophic levels based on both quantitative and qualitative data on diet items, mirroring the approach employed in estimating species’ trophic levels within the FishBase database.

### Reconstruction of food webs time series

To reconstruct the food webs for each fish community at each time step, we modeled the probability that a focal species preys on a candidate species with which it co-occurs, applying a trait-matching function based on the niche model for food web structure ^75^, where the log body size of the predator determines its optimum and the range of its trophic niche, while log prey size determines its position in the predator niche ^30^.Thus, with available data on observed predator-prey interactions and their body size, the model can be calibrated to define a predation window defining the body size range within which a predator species of a specific size can feed ^30,76^. Ultimately, the model indicates whether the focal consumer species preys on a candidate resource species.

To calibrate and assess the performance of the model, we used a data subset of predator-prey interactions derived from the data compilation on species diets (see data collection and selection, species diets). This data subset encompasses 23,834 predator-prey interactions at the species level, involving 1,396 predator species and 2,515 prey species. These interactions cover diverse locations in both freshwater and marine environments, capturing a broad spectrum of environmental conditions (see Supplementary Data).

To select the model used in the reconstruction of the food webs, we evaluated multiple models with different windows of predation (quantiles 0.01-0.99, 0.02-0.98, 0.03-0.97, 0.04-0.96, 0.05-0.95) in different fish groups (marine and freshwater). For each model, we split the predator-prey dataset into two parts, 70% of the dataset was randomly chosen to calibrate the model, while the remaining 30% was used to evaluate it. We calculated the Boyce index to evaluate the quality of the models and select those with the highest values (Marine fish 0.05-0.95, mean Boyce index = 0.56, s.d. = 0.09, n = 999; Freshwater fish 0.03-0.97, mean Boyce index = 0.53, s.d. = 0.08, n = 999 Extended data Fig. 2). The Boyce index varies from −1 to +1 where a positive value indicates a model whose predictions are consistent with the presence of interactions in the evaluation dataset, values close to zero mean that the model is not different from a random interaction, while values values close to −1 indicate an incorrect model which predicts unsuitable interactions ^77,78^. This cross-validation procedure was repeated 999 times. After identifying the most performant models, we applied them separately to the 2,090 marine and 822 freshwater species present in the assemblages time series using their respective body size data (refer to the Body size section in Data collection and selection). This enabled us to identify all potential predator-prey interactions that may occur among all the species in the database, and subsequently allocate them only to the specific pair of species (predator-prey) that co-occur in a particular sampling. Because of the size-base nature of our model, it can misidentify trophic interactions forbidden in nature (e.g., large herbivorous fish feeding on smaller fish). To correct this potential issue, we removed all predator interactions for fishes whose documented diet was not classified as piscivorous.

Modeling predator-prey interactions based on species traits is a common approach for predicting interactions within food webs across different scenarios of species composition. This includes the reconstruction of historical food webs based on past species composition and the generation of current food webs in regions where direct recordings are lacking (e.g. ^29,31,79,80^). Besides, this method helps to overcome sampling uncertainty inherent in traditional methods, by identifying likely interactions that may not have been detected and filling gaps in undersampled ecological networks. Ecologists typically have a limited understanding of food webs, as even intensive sampling efforts often fall short in fully capturing interactions ^81,82^. The difficulty lies in detecting interactions, which are often infrequent or occur between rare species ^81–83^.

Recognizing this, there has been a growing effort over the past decade to construct models that forecast unobserved links within ecological networks ^84,85^. Our study builds on these previous efforts, aiming to capture all potential interactions among the set of co-occurring fish species even if not yet documented in the literature, driven by the imperative to better understand ecological networks and their transformations resulting from human-induced effects in biodiversity.

Although trait-based inferences can be advantageous in reconstructing predator-prey interactions, relying solely on traits also comes with some limitations. This is because a singular trait value per species overlooks intraspecific trait variation, such as potential variations between populations of the same species in different environments ^86^, which could potentially lead to an underestimation of niche breadth and species interactions ^87^. The selection of trait and interaction records from global databases, particularly for rare or poorly studied species, could also represent another crucial source of uncertainty and bias as these records are more likely to capture out-of-the-range observations, ultimately affecting model predictions in (e.g. ^88^).

### Dataset description

For the analysis presented here, the database comprises 100 replicates (or iterations) of 15,029 food web time series reconstructed for rarefied fish assemblages data from 103 different studies or monitoring programs, and including 2,844 species. Specifically, 12,467 of these time series focus on marine environments, while 2,562 focus on freshwater environments. To ensure a sufficiently large network size for metric measurement, all time series adhere to a minimum rule of having at least five species across the sampling period. Geographically, the data exhibit a notable skew towards Europe, North America, and Australasia, as illustrated in the extended Figure 1. The time series in the database span a mean duration of 16.84 (± 10.75) years, with a corresponding mean of 6.31 (± 5.56) sampling years. For accessibility, all utilized data are available in the code repository listed in the Supplementary Data.

### Calculation of taxonomic diversity, body size and food web metrics

Within each time series, we calculated several metrics describing different biodiversity facets:

#### Taxonomic diversity

- Temporal dissimilarity in species composition: Evaluated through pairwise Jaccard dissimilarity between the first year and subsequent years in the time series.
- Species richness: Determined as the number of species/nodes in a given year in a time series.

#### Body size

- Mean body size: average body size of species in a given year in a time series.

#### Food webs

Within the food webs we calculated different metrics related with the network topology, species’ role and food web functioning.

#### Topological

- Connectance: The proportion of realized interactions out of all possible bipartite interactions, expressed as ratio between number of links and square of the number of species.
- Modularity: Describes the presence of aggregated sets of interacting species, indicating the extent to which interactions occur more frequently within modules than between modules.

#### Species’ role

- Generality: represents the mean number of prey taxa consumed per predator.
- Prey vulnerability: indicates the mean number of predators consuming each prey taxon.
- Mean trophic similarity: Calculated as the average trophic similarities between all species in a food web. Trophic similarity between two species is defined by the number of common prey and predators divided by their total number of preys and predators.

#### Functioning

- Trophic group proportions: the proportion of species within a specific trophic group (i.e. top predators, mesopredators, omnivorous, primary consumers) relative to the total number of species within the food web.

### Statistical analysis - Models of trends

We employed mixed-effects models to examine temporal variations in all biodiversity metrics, encompassing taxonomic diversity, body size, and food web metrics. In the fixed model structure, the selected biodiversity metric served as the response variable, with Year (mean-centered) as the sole fixed independent variable. Additionally, Year was included as a random slope, varying across studies and spatial units, which represent the smallest reported sampling units. To address the non-independence of the time series, spatial units were nested within the original studies from which they were derived. We utilized poisson, normal, and binomial error structures to model biodiversity changes based on the distribution of the response variable (i.e., biodiversity metric). All statistical models were implemented within a frequentist framework using the ‘glmmTMB’ package ^86^ in R (v4.2.1; ^89^). The overarching model structure in ‘glmmTMB’ annotation is as follows:

\# gaussian family function glmmTMB(biodiversity metric ∼ Year_Cent + (Year_Cent|Study_ID_All/Spatial_Uni_ID), data

= data_Mod_TL_BS2, family = gaussian) # poisson family function glmmTMB(biodiversity metric ∼ Year_Cent + (Year_Cent|Study_ID_All/Spatial_Uni_ID), data

= data_Mod_TL_BS2, family = poisson) # binomial family function glmmTMB(cbind(Number of species in a trophic level, Total species in the assemblage) ∼ Year_Cent + (Year_Cent|Study_ID_All/Spatial_Uni_ID), data = data_Mod_TL_BS2, family = binomial)

Each biodiversity metric’s model was fitted to the 100 iterations of the time series datasets to account for potential variations in species composition resulting from standardizing sample effort using sample-based rarefaction (see Methods, data collection, and selection, assemblage time-series data). Subsequently, we extracted the slope from each model fit and combined them to draw inferences by comparing the mean coefficient and the confidence intervals across 100 iterations, accounting for the uncertainty in the assemblage time series.

## Code and data availability

All the code and data used for the analyses will be made available upon acceptance. The raw data on assemblage time series are available at https://biotime.st-andrews.ac.uk/download.php (ref. ^37^) and https://idata.idiv.de/ddm/Data/ShowData/1873?version=12 (ref. ^36^) in so far as they are not openly available elsewhere.

## Acknowledgements

We thank C. Schmidt for the illustration work in the figures and S. Blowes for advice on the model used. Funding: German Centre for Integrative Biodiversity Research (iDiv) Halle-Jena-Leipzig (German Research Foundation FZT-118 grant 202548816).

## Author contributions

J.C.Q. conceived the initial idea. J.MC, U.B., M.D. and L.C. further developed and designed the research concepts. J.C.Q. and J.H.P collected the data. J.C.Q. performed the statistical analysis and created the figures. J.C.Q. wrote the initial manuscript draft, and J.MC, U.B., M.D., L.C., P.A.T., and X.G. provided comments on the manuscript.

---

**Extended Data Fig. 1.**
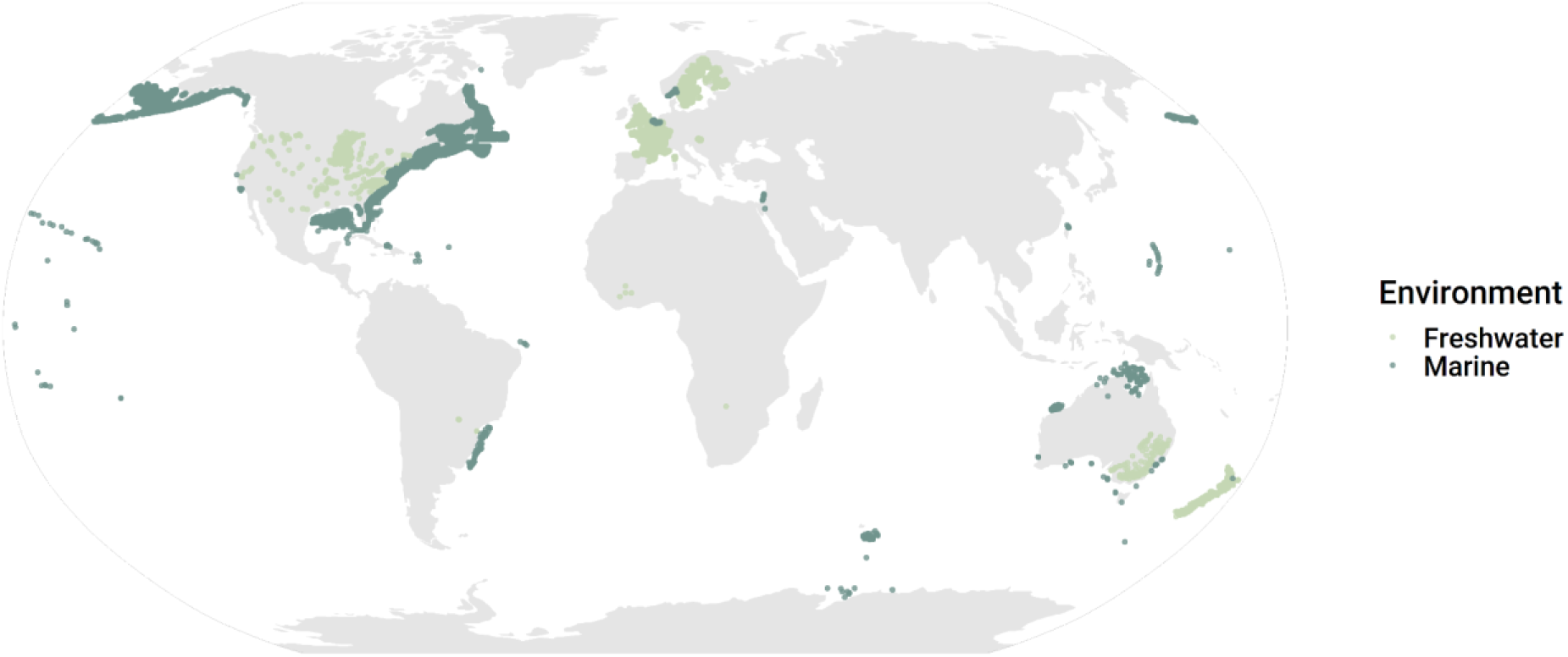
Bivariate map depicting the spatial distribution of time series at the spatial unit level. Each dot denotes a fish-assemblage time series with a reconstructed food web. Dark green dots denote fish assemblages in marine environments, while light green denotes freshwater assemblages.

**Extended Data Fig. 2.**
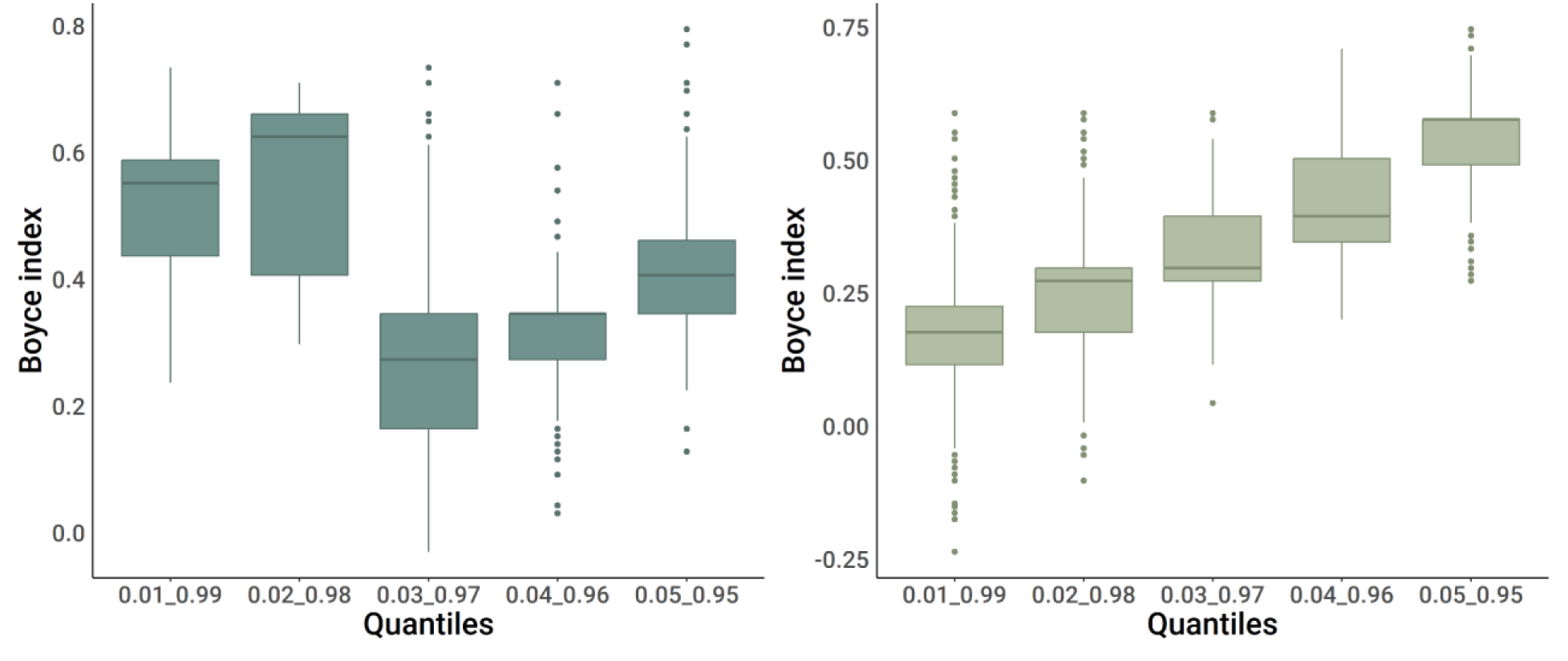
Assessment of niche model calibration process. The niche model was fine-tuned using an independent dataset of predator-prey interactions, employing various predation windows (quantiles) for fish species found in marine (dark green) and freshwater environments (light green). Models exhibiting superior performance, as determined by the Boyce index (Marine fish 0.05-0.95, mean Boyce index = 0.56, s.d. = 0.09, n = 999; Freshwater fish 0.03-0.97, mean Boyce index = 0.53, s.d. = 0.08, n = 999), were chosen to reconstruct trophic interactions within the fish assemblages.

**Extended Data Fig. 3.**
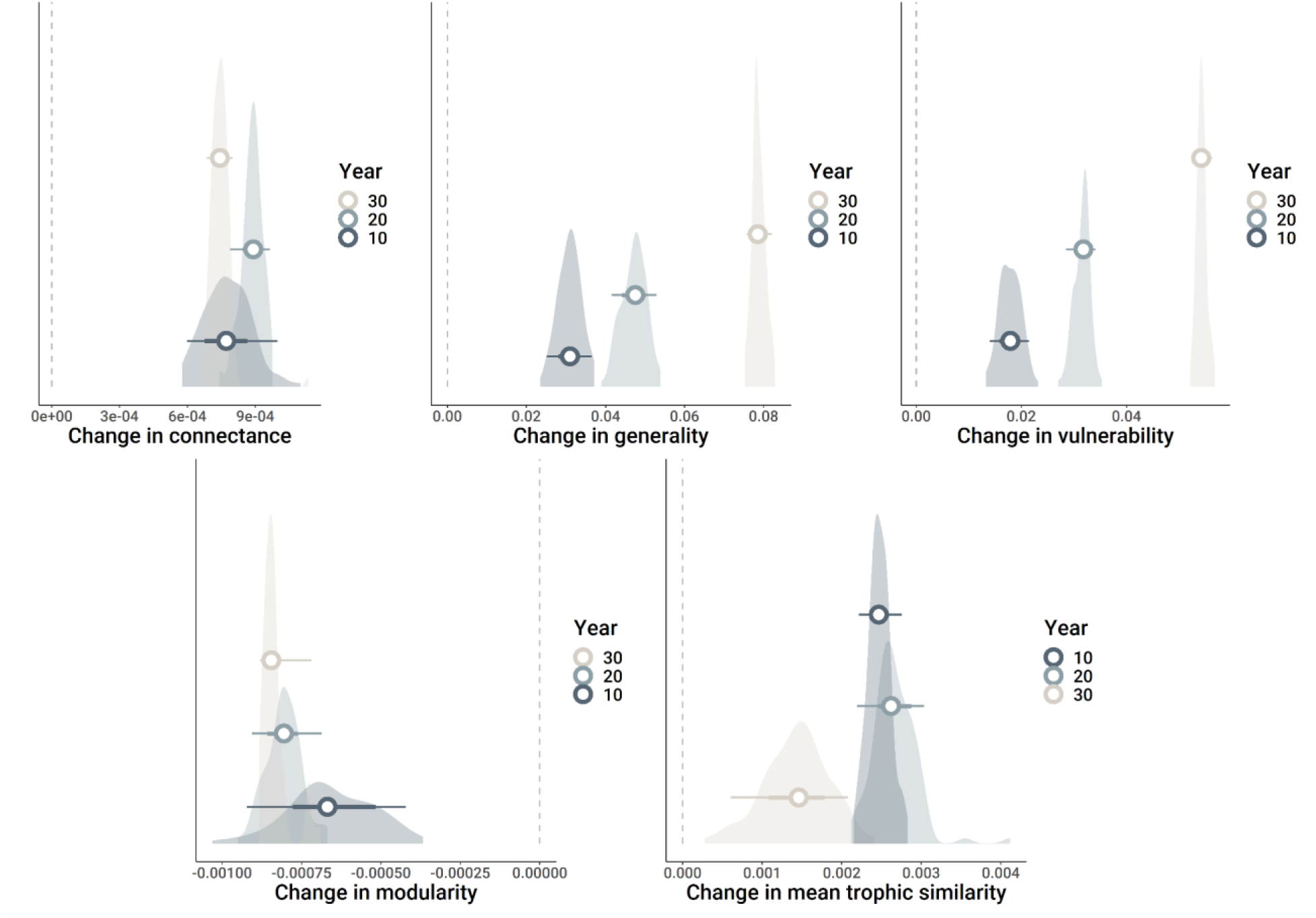
Changes in food web topology metrics and trophic similarity in time series spanning at least 10, 20 and 30 years. The density plots display the distribution of slopes for changes in connectance, generality, prey vulnerability (predation pressure), modularity, and trophic similarity. The horizontal bars with error bars denote the mean and the 50% and 95% confidence intervals (CIs) of the mean estimates (depicted by white circles).

**Extended Data Fig. 4.**
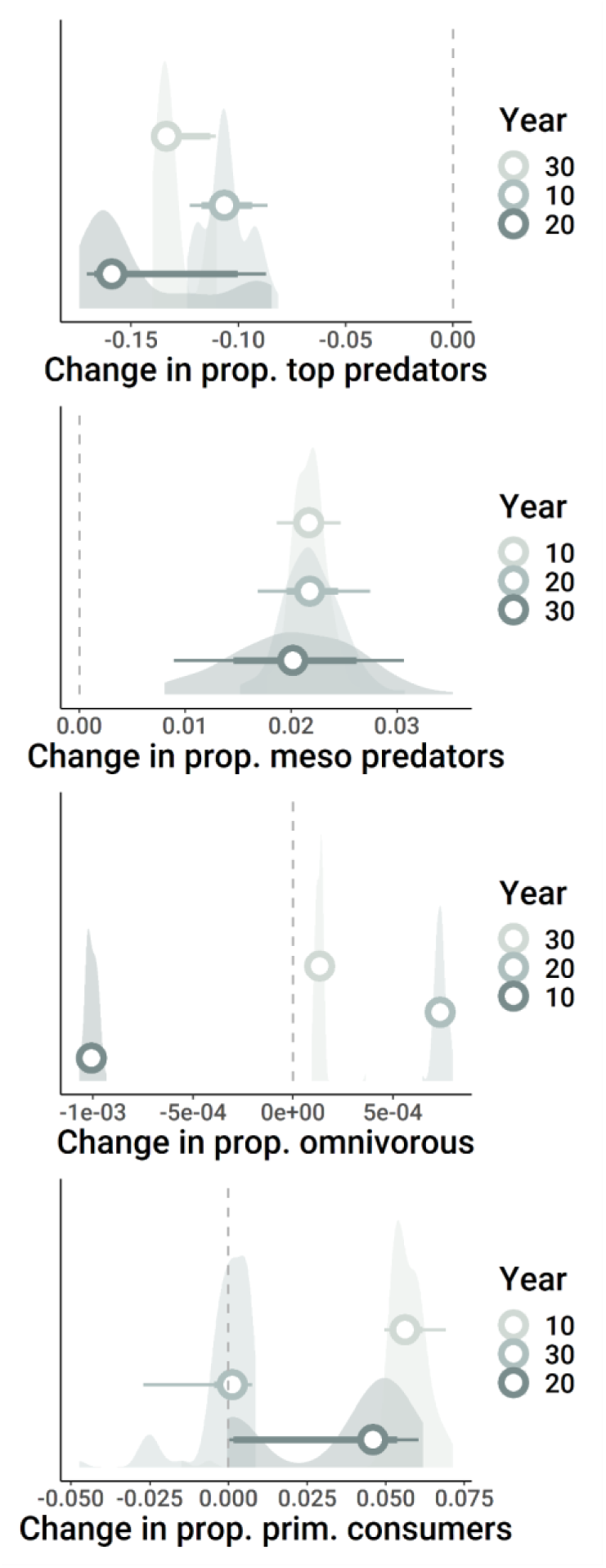
Changes in proportion of trophic groups in time series spanning at least 10, 20 and 30 years. The density plots display the distribution of slopes for changes in proportion within the assemblages of top predators, mesopredators, omnivorous, and primary consumers. The horizontal bars with error bars denote the mean and the 50% and 95% confidence intervals (CIs) of the mean estimates (depicted by white circles).

**Extended Data Fig. 5.**
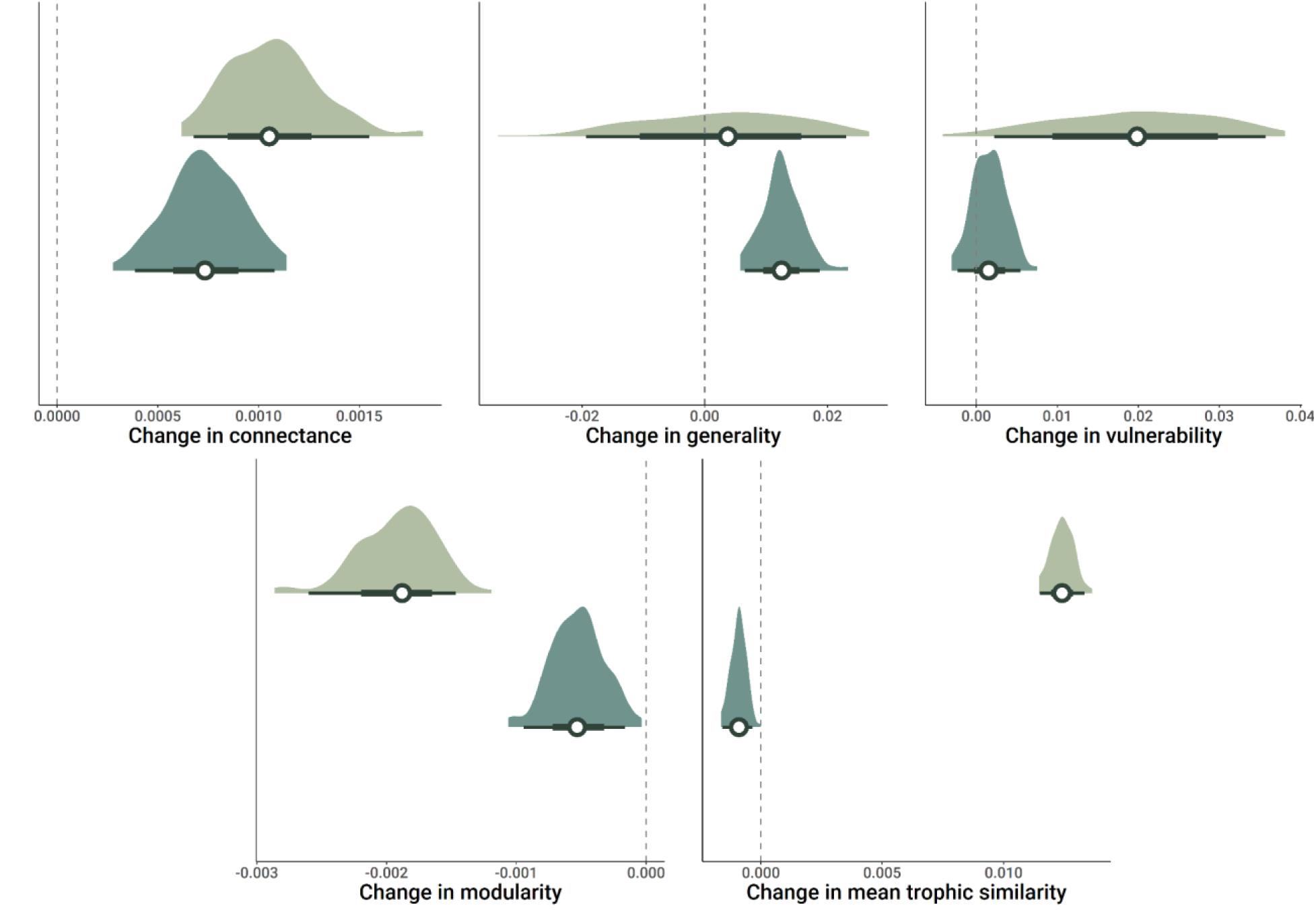
Changes in food web topology metrics and trophic similarity in marine (dark green) and freshwater environments (light green). The density plots display the distribution of slopes for changes in connectance, generality, vulnerability (predation pressure), Modularity, and trophic similarity across the 100 iterations of the rarefied time series. The horizontal bars with error bars denote the mean and the 50% and 95% confidence intervals (CIs) of the mean estimates (depicted by white circles).

**Extended Data Fig. 6.**
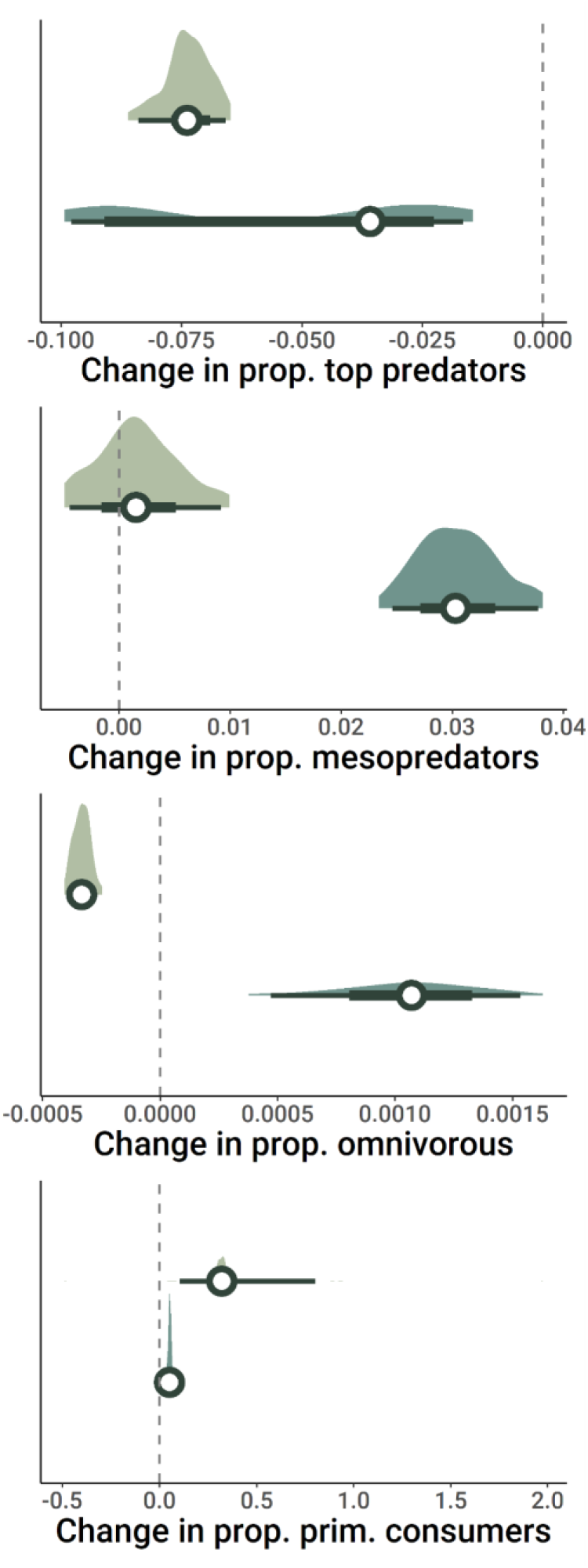
Changes in proportion of trophic groups between marine (dark green) and freshwater environments (light green). The density plots display the distribution of slopes for changes in proportion within the assemblages of top predators, mesopredators, omnivorous, and primary consumers. The horizontal bars with error bars denote the mean and the 50% and 95% confidence intervals (CIs) of the mean estimates (depicted by white circles).

